# Chemotaxis and topotaxis add vectorially for amoeboid cell migration

**DOI:** 10.1101/735779

**Authors:** Joeri A. J. Wondergem, Maria Mytiliniou, Falko C. H. de Wit, Thom G. A. Reuvers, David Holcman, Doris Heinrich

## Abstract

Cells encounter a wide variety of physical and chemical cues when navigating their native environments. However, their response to multiple simultaneous cues is not yet clear. In particular, the influence of topography, in the presence of a chemotactic gradient, on their migratory behavior is understudied. Here, we investigate the effects of topographical guidance on highly motile amoeboid cell migration (topotaxis) generated by asymmetrically placed micropillars. The micropillar field allows for an additional, natural chemotactic gradient in two different directions, thereby revealing the relevance of topotaxis in the presence of cell migration directed by chemical gradients (chemotaxis). Interestingly, we found that the topotactic drift generated by the pillar field is conserved during chemotaxis. We show that the drifts generated by both these cues add up linearly. A coarse-grained analysis as a function of pillar spacing subsequently revealed that the strength and direction of the topotactic drift is determined by (i) the pore size, (ii) space between pores, and (iii) the effective diffusion constant of the cells. Finally, we argue that topotaxis must be conserved during chemotaxis, as it is an emergent property of both the asymmetric properties of the pillar field and the inherent stochasticity of (biased) amoeboid migration.

## Introduction

Directed, single cell amoeboid migration is driven by external guidance cues, such as chemical, electrical, temperature, stiffness, and topographical gradients^1–6^. Natural cell environments often exhibit several such cues simultaneously. In the human body, processes occurring in multi-cue environments include immune response^7,8^, cancer metastasis^9,10^, and tissue regeneration^11^. As of yet, it is unclear how external guidance cues relate to each other for various cell types and environments. Cells may ignore certain stimuli in favor of other cues or different cues might add up in affecting cell movement. In this paper, we focus on the effects of long-range topographic guidance on cell migration and investigate whether observed topographical guidance is conserved during chemotaxis.

The physical structure of the micro-environment around the cell, the 3D-topography, can act as a stimulus modulating cell direction and speed during migration^6^. *In vitro* topographical structures are typically made up out of channels^12–15^, ratchets^16–18^, grooves^19–21^, pillars^22–24^, curves ^25^, or areas with increased alignment of extracellular matrix fibers^14,26–28^. These *topotaxis* assays rely on topographical asymmetries, and exploit differences in slope, confinement or alignment to influence cell migration. The aforementioned studies focus largely on asymmetries much smaller than the cell body, at the length scale of extracellular matrix fibers, to investigate topographic guidance effects.

Instead, we investigate long-range topographical guidance that arises from asymmetrically distributed physical obstacles of cell size. Such environments are encountered by highly motile cells navigating entire tissues, like macrophages during immune response and metastasizing amoeboid sarcoma cells. For example, recent studies of neutrophil movement inside Zebra fish larvae map migratory patterns across the fish tail following a chemotactic response^29,30^. The migration of these cells is effectively a (biased) random walk through a crowded environment. However, the role of possible long-range topotactic effects, emerging from differences in tissue organization, as the cell travels long distances is unclear.

To explore the influence of a physical gradient in crowded environments, we engineered anisotropic micro-pillar fields and recorded cell migration through these fields. Because cells navigating tissue are often exposed to chemotactic processes, we presented the cells migrating in these pillar fields with an additional cue and gauged if any topotactic effects are conserved. The potential interplay between topographical and chemical guidance cues has not been studied extensively (cf. Comelles et al. on topo-versus haptotaxis^17^). Yet, such combined assays provide an excellent platform to gain insight into how multiple cues affect cell migration simultaneously. It could even be argued that the strength of measured topotactic guidance effects can only be gauged in the context of other stimuli, most notably chemical gradients.

We used *Dictyostelium discoideum (D. discoideum)* as a model organism^31^ to address these questions. When starved, these cells show a strong chemotactic response to cyclic-adenosine monophosphate (cAMP) gradients^32^. During migration in this starved state, cells repeat a process of pseudopod splitting and elongation^3,33,34^. This amoeboid migration is evolutionarily conserved for many other cell-types, like metastasizing cancer cells^35^ and leukocytes^36^. Trajectories of such cells, and *D. discoideum*, display a mix of random motion (pseudopod splitting) and persistence (elongation) at very short time scales, but evolve into a random walk over longer periods of time when devoid of a clear chemical signal^37^. These persistent random walks become biased when external guidance cues are present, resulting in an effective drift. Aside from chemical gradients, *D. discoideum* migration can be influenced topographically^38^. Thus, embedding the amoeba in asymmetric pillar fields, overlaid with a chemical gradient, yields a topo-chemical multi-cue environment capable of studying the significance of topotaxis for highly motile amoeboid movement.

As mentioned before, cell migration through a field of pillars, or through a tissue topography, is reminiscent of a random walk through a crowded environment. Therefore, in the second part of this paper, we explore how long-range topographical guidance can arise from just the stochasticity of cell trajectories. To that end, we model cell movement through the pillar field as a jump process between different pillar spacings and effectively coarse-grain the dynamics of the system to the scale of domains larger than the cell size. We show that random movement in an asymmetric pillar field must result in topotaxis. This means that every multi-cue system is influenced by the topographic asymmetries present, as there is an inherently random component in cellular response to any stimulus. Therefore, we argue that topotactic effects, caused by physical asymmetries, form a fundamental component of navigation for *in vivo* systems, in which many different cues are at play.

## Results

### Experimental assay combining the effect of topographic and chemical cues

We studied amoeboid cell movement inside a micropillar field of periodically increased spacing (fig. **1**) using the model organism *D. discoideum* in the starved state. As we will show, such an asymmetric pillar field functions as a topotactic cue. Subsequently, both the pillars and cells were embedded in a microfluidic device enabling an additional chemotactic gradient by controlled diffusion (**M&M**) of cyclic-adenosine monophosphate (cAMP), a well-studied chemical attractor for *D. discoideum*^3,39^. By adding the second gradient, we created an experimental assay with two competing stimuli for cell motility: an asymmetric topography and a chemical gradient in solution. First, cell movement was studied in the pillar field without a chemotactic gradient (fig. **1a**). Then, to investigate if topotaxis (T) is conserved for chemotaxing cells, we introduced cAMP gradients aligned (A) and opposed (O) to the topographical guidance cue in either the negative (fig. **1b**) or positive (fig. **1c**) *x*-direction. These two configurations ensure competitive or synergistic setups for both external stimuli.

**Figure 1:**
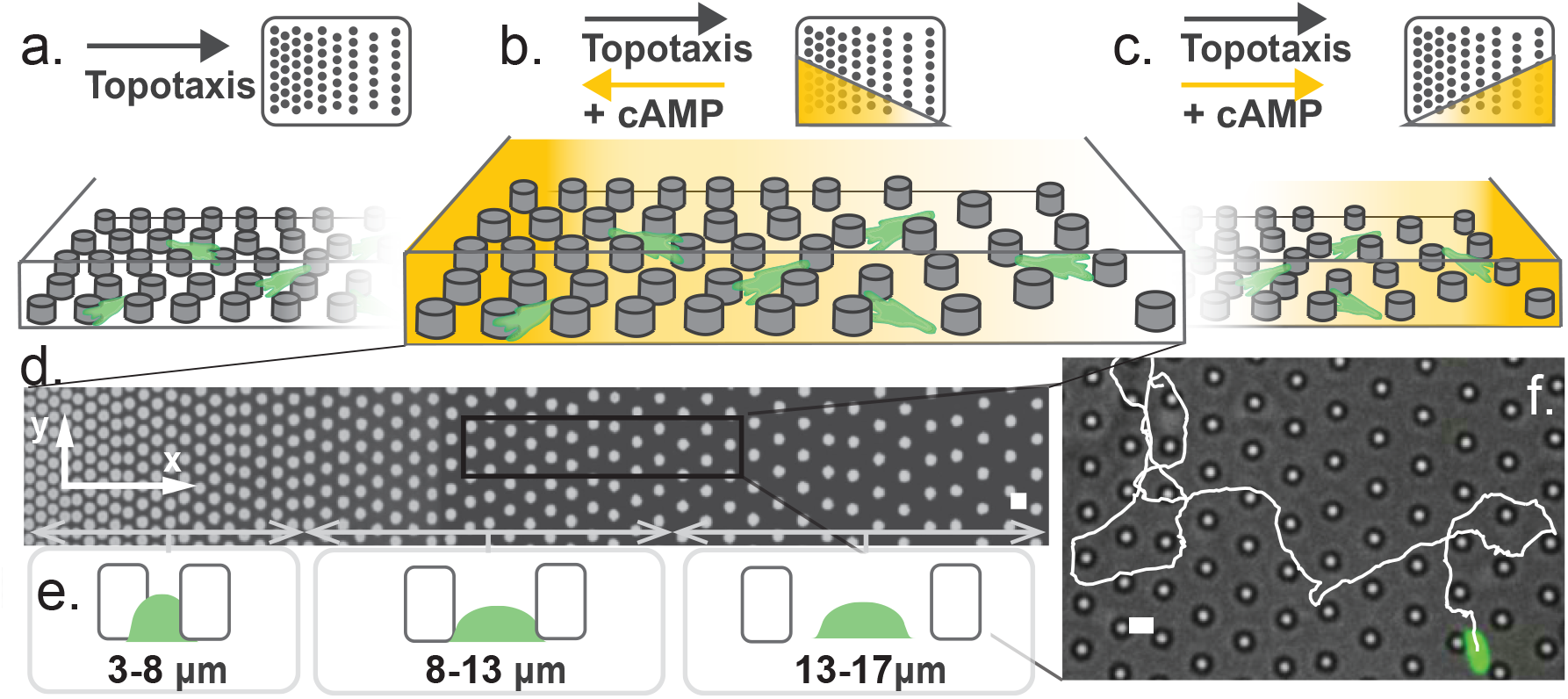
The experimental assay. **a.** Cells are seeded on a PDMS micropillar array with varying lattice spacing, generating a topotactic gradient (T). **b.-c.** An additional chemotactic gradient (cAMP, yellow) is applied, opposed (O) or aligned (A) to the topotactic gradient. **d.** Fluorescent image of the full micropillar array, overlaid with the orientation of the axis chosen. Characteristic spacing (wall-to-wall) between the pillars below the image. **e.** Three characteristic spacings of the pillar array: smaller than the cell diameter, approximately the cell diameter and larger than the cell. **f.** Example trajectory of starved *D. discoideum* (green) inside the micropillar array (bright field image). Scale bars are 10 *µm*.

Figure **1d** shows a top view of the pillar field. The pillars are fabricated from the transparent polymer polydimethylsiloxane (PDMS) and assumed chemically homogenous (**M&M**). The height and diameter of the pillars is the same (8 µm) and comparable to the typical size of a *D. discoideum* cell, which varies between 8-15 µm ^40^. This ratio between pillar height and cell size is sufficient to keep the motion of starved *D. discoideum* two-dimensional, meaning the pillars act as obstacles, as observed in previous studies^38,41^. Cells fully mounting the pillars are rarely observed and excluded from the data. The space between pillars increases from 3 to 17 µm in the *x*-direction resulting in pillar center-to-center distances between 11-25 µm.

The pillar field (figure **1d**) is arranged in a trigonal lattice to ensure cells encounter pillars during migration in the direction of the spatial gradient (+*x*-direction) and chemotactic gradients (±*x*-direction), thereby excluding the possibility of cells simply migrating along pillar alleys. The pillar array spacing increases by 1 µm after every fourth pillar (in +*x*). By not increasing the distance between every subsequent pillar, cells more likely reorient at least once within each spacing and explore it, as their migration trajectories exhibit a persistence length between 20-40 µm on a flat surface (**S1**). Overall, the available space between pillars spans three different length scales as compared to the cell diameter (fig. **1e**): a squeezed state (3-9 µm), a relaxed state with continuous pillar contact (9-13 µm), and a state where the cell periodically loses contact with pillars (13-17 µm), travelling from pillar to pillar. Still, cells move smoothly through each spacing chosen. The pillar field exhibits a topographic asymmetry that is much larger than the cell body and therefore functions, in effect, as a series of tissue pores of increasing spacing.

### Cell migration inside an asymmetric micro-pillar array

To quantify cell migration in the topotactic gradient, we analyzed cell center-of-mass trajectories. Although this approach ignores individual intracellular dynamics, it enables multi-cell tracking over a long period of time, which, in turn, allows for statistical analyses, to determine the overall effect of external gradient fields by quantifying average drift^42^. The acquired cell trajectories within the anisotropic pillar field are plotted centered in figure **2a**. Figure **1f** (**Movie 1**) shows one of these trajectories projected onto the pillar field. Cell motion in an anisotropic pillar field produces similar trajectories as compared to those on a flat surface (**Movie 2**), but there is a general preference to the more spacious side of the pillar array and this preference is a topotactic (T) effect. Upcoming, we will show that this effect is dependent on both the increase in surface area available to each cell and the space between pillars also available for the cells. Furthermore, when the topotactic cue is tested versus the chemotactic cue, topotaxis is conserved. In the aligned configuration (A) cells are clearly drawn towards the source of cAMP (fig. **2c**) more, than when topographical and chemical cues oppose (O, fig. **2b**). The asymmetric pillar field decreases the directional response of cells to a cAMP source.

**Figure 2:**
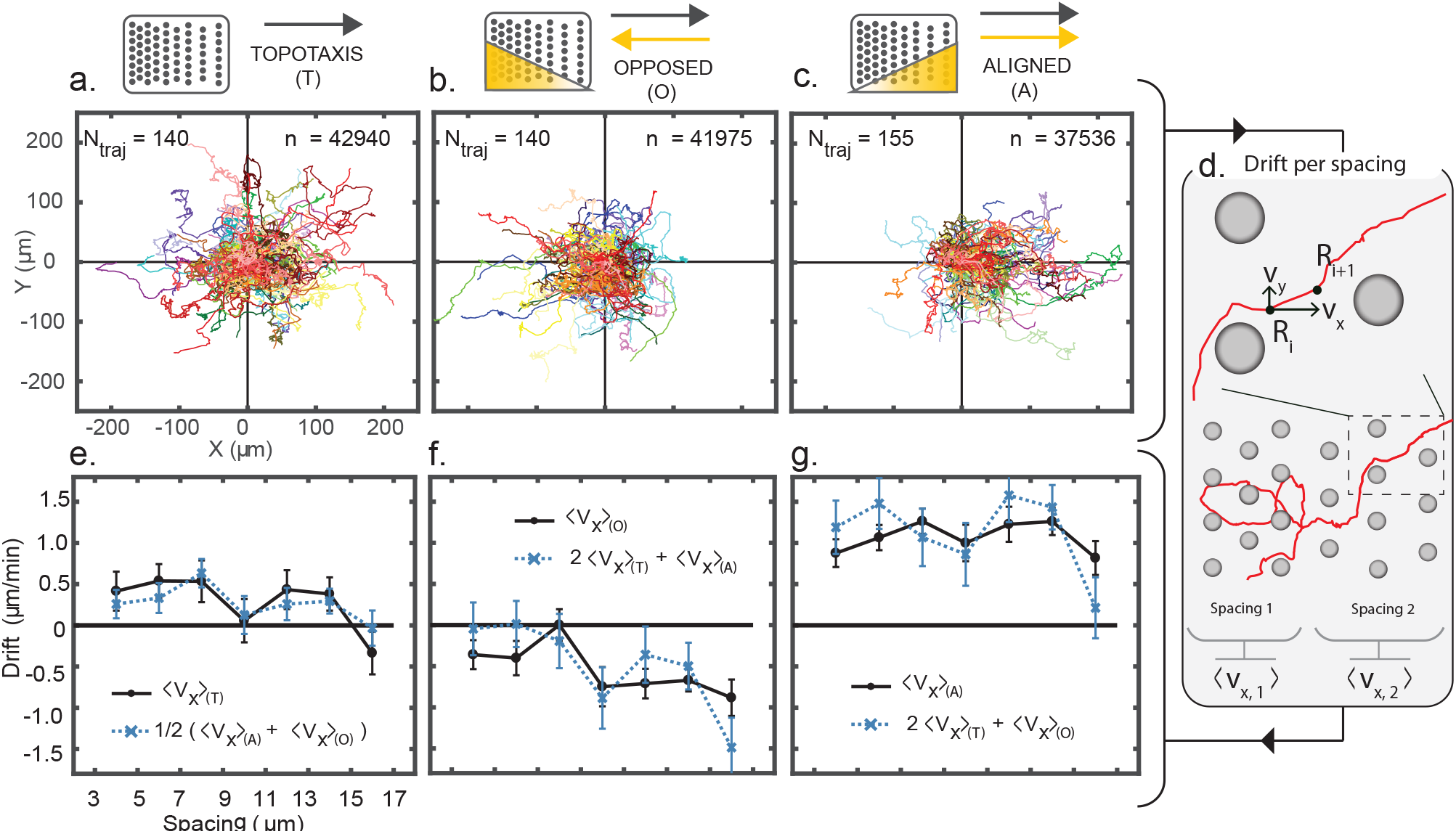
Topotaxis is conserved during chemotaxis. **a.-c.** Cell trajectories for the three experiments performed: topotaxis, opposed and aligned. Cells migrate less towards the chemical attractor than when both cues are aligned. **d.** Schematic of calculating the drift. All displacements in the x-direction are averaged per pillar spacing and per experiment. **e.** Drift measured in the direction of the topotactic gradient (black line) as a function of the interpillar distance. Error bars are 95% confidence intervals. Topotactic drift is similar to the ‘reconstructed’ topotactic drift (blue line), found by adding the drifts of the other two experiments. This indicates topotaxis is conserved during chemotaxis. **f.-g.** Drift measured in the opposed and aligned experiments (black). The data closely follows the ‘reconstructed’ drifts from the other experiments (blue), also indicating topotaxis is conserved.

To quantify the difference, the drift 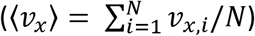 was calculated as a function of interpillar spacing (fig. **2d**, **M&M** and **S2**) for all cell migration trajectories and shown in figures **2e-g** (black lines) for all experiments. In the aligned experiment, the drift is generally 〈*ν*_x,A_〉 ≈ 1 µmmin^−1^, whereas in the opposed and in the topotaxis experiments the drifts measure between −〈*ν*_x,O_〉 ≈ 0.5 − 1 µmmin^−1^ and 〈*ν*_x,T_〉 ≈ 0 − 0.5 µmmin^−1^ respectively. These drifts not only show that the topotactic gradient influences the chemotactic response when the cues are opposed, but, interestingly, they indicate linear summation, as 2 ⋅ 〈*ν*_x,T_〉 ≈ 〈*ν*_x,A_〉 + 〈*ν*_x,O_〉. To be the case, the topotactic drift 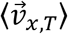 should be similar to the ‘reconstructed topotactic drift’, found by adding the drifts of the other two experiments (A&O), which is defined as 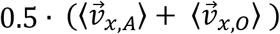. Figure **2e** (blue line) shows that this is indeed the case. When the procedure of linear addition is repeated for the other two experiments, the ‘reconstructed’ aligned and opposed drifts, respectively 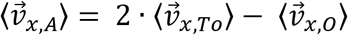 and 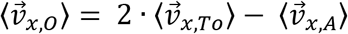 (fig. **2f-g**, blue line), also follow the measurements closely. To conclude, these findings suggest that the topotactic response can be independently deduced from the experiments with the double gradients, showing that the topotactic and chemotactic drifts add up linearly.

### Coarse-grained analysis: cells transitioning between different pillar spacings

Cell movement in the anisotropic pillar field with respect to the spatial gradient is effectively a series of transitions between bands (in direction ±*y*) of equal pillar spacing (fig. **3a-c**, **M&M**). A cell at a certain spacing (*x*_*k*_) can either move towards a more spacious (*x*_*k*−1_, fig. **3a-c**, orange) or less spacious pillar spacing (*x*_*k*+1_, fig. **3a-c**, green). By recording all cell transitions between different spacings, and identifying them as a function of the pillar gradient, we computed the transition probability towards the more spacious side (*P*_spacious_) and less spacious side (*P*_narrow_), and subsequently their difference (Δ*P*) (fig. **3c** and **M&M**). Figures **3a-c** (**Movie 3**) shows this analysis for a single cell migration trajectory. The transition probabilities are displayed in a histogram exhibiting the net directional preference of cell migration with respect to the spatial inter-pillar gradient (fig. **3c**). By coarse-graining the whole lattice spacing, as opposed to a time-dependent (Δ*t* = 8 − 10 s) direct drift calculation (〈*ν*_x_〉), we found that the noise in the overall directional preference was reduced.

**Figure 3:**
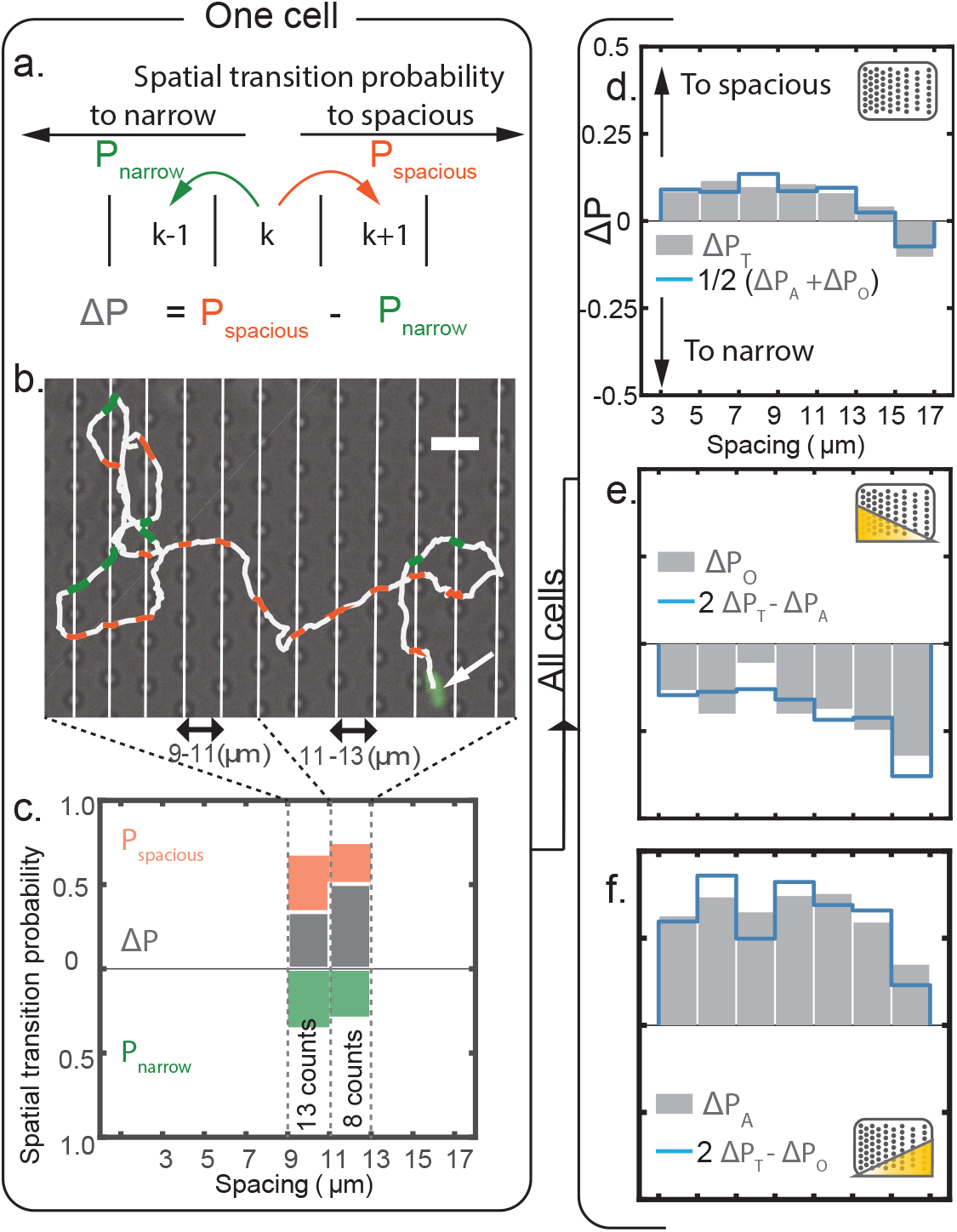
Spatial transition probability histograms for all three experiments. **a.-b.** Micropillar array is computationally divided to track the dynamics of the cell (arrow) as a function of the spacing between pillars. Each time a cell transitions to a new domain, with respect to the direction of the gradient, a transition is recorded to either the more spacious (orange) or narrower (green) part of the pillar field. Scale bar is 15*µm*. **c.** A spatial transition probability histogram can be made using the recorded transitions, seperating the probability to transition to narrower (*P*_*orange*_) or to more spacious (*P*_*green*_) at each pillar spacing. **d.** Histogram of all transitions measured for the topotaxis experiment, showing that cells migrate to areas with bigger pillar spacing (grey). The ‘reconstructed’ ∆*P* (blue) shows that the difference between the opposed and aligned experimenst is exactly the topotactic cue. **e.-f.** Histograms with the topotactic and chemotactic effects opposed and aligned, respectively

The transition histogram for all cell migration trajectories in the anisotropic pillar lattice (fig. **3d**) shows that cells are topographically guided towards the spacious region of the pillar array for spacings 3 − 15µm. The difference in cell transition probabilities ranges between Δ*P*_T_ = 0.10 − 0.15. However, when pillars are spaced further apart than the cell size (15 − 17µm), cells prefer migration to the opposite direction (Δ*P*_*T*_ = 0.02 − 0.09). This result indicates that cells have an affinity to pillars spaced equal to their cell diameter. This result is reproduced on another pillar field with a steeper inter-pillar gradient (fig. **S4**). When the chemical and topographical gradients are aligned (fig. **3f**), cells transition more readily Δ*P*_*A*_ = 0.23 − 0.43 towards the source of cAMP compared to when the gradients oppose (fig. **3e**) Δ*P*_O_ = 0.15 − 0.24. Again, this indicates that the topotactic gradient influences chemotactic response.

To study the combinatorial response, we analyzed the ‘vectorial sum’ of the transition probabilities. The topotactic cue can be distinctly reconstructed from the aligned and opposed experiments (Δ*P*_*T*_ = 0.5 ⋅ (Δ*P*_*A*_ + Δ*P*_*O*_), fig. **3d**, blue line). Similarly, twice the difference between the topotaxis and either aligned or opposed gradient experiments should generate the other: Δ*P*_*A*_ = 2 ⋅ Δ*P*_*T*_ − Δ*P*_*O*_ and Δ*P*_*O*_ = 2 ⋅ Δ*P*_*T*_ − Δ*P*_*A*_. The histograms shown in figure **3e**–**f** (blue lines) confirm that the aligned and opposed response can be reconstructed from the opposite experiments respectively. The similarity between both the measured signals (black lines) and the reconstructed signals (blue lines), for all three experiments, shows that the same topographical guidance is indeed measured in the experiments with and without a chemical gradient. These results therefore confirm that this topographical cue is conserved during cellular chemotaxis. Moreover, the topotactic and chemotactic drifts add linearly.

Next, we develop a model to explain the migratory drift generated by the spatial gradient in the pillar field. The cornerstone of this model is to coarse-grain cell movement throughout the pillar lattice as a series of escapes of a Brownian particle from domains delimited by pillars. Using both the physical model and the experimental results, we will explore the implications of spatial gradients for cell migration in multi-cue environments.

### Modelling topotactic drift in a pillar lattice

Cell movement inside the pillar field is, in essence, a series of escapes from quadrilateral domains delimited by pillars (fig. **4a**). The average time to escape from an arbitrary point inside a predefined domain to the boundary of the domain is the mean first passage time (MFPT). We will empirically estimate the MFPT of trajectories as the duration between the entry and exit into and out of a quadrilateral pillar domain. To measure these times, we first identify the edges of each domain by linking all the pillar centers (**M&M**) as shown in figure **4a** (grey lines). Each escape from domain to domain, is then registered (fig. **4a**, green, **Movie 4**) and yields a collection of first passage events (*n* = 4002) for each pillar spacing. The passage time distributions for different pillar spacing are shown in figure **4b** (and **S4**). Note that for each spacing the MFPTs are well fit by a single exponential decay. This suggests that the escape from each domain is a Poissonian process and it is conserved over the entire pillar field.

**Figure 4:**
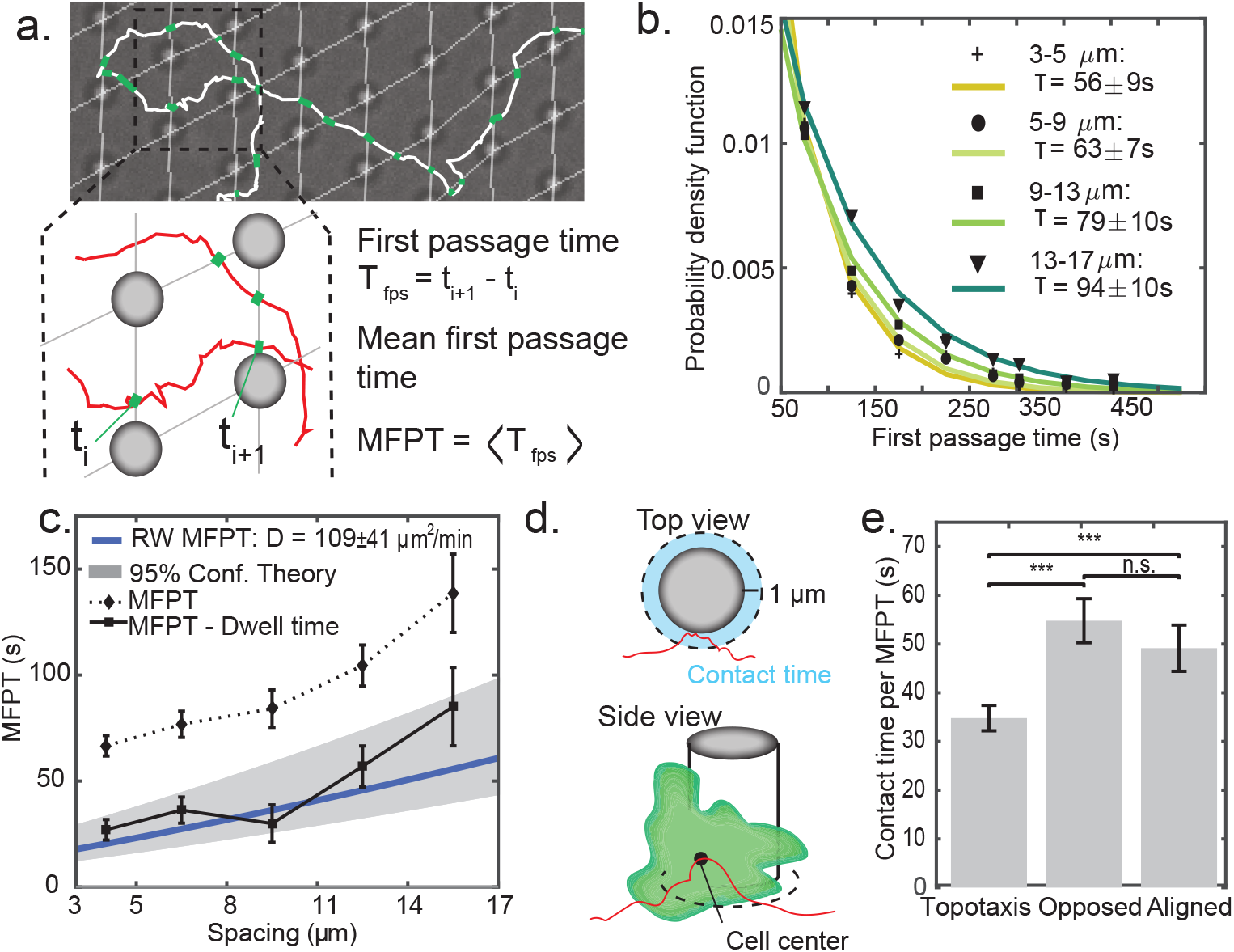
Cell centers escape as a Brownian particle from pillar domains. **a.** Example trajectory of a cell moving through pillar domains. Transitions into a new domain are recorded (green) and the passage time between entry and exit computed. **b.** Probability density function of the first passage times (*T*_*fps*_) measured for different spacing. The graphs suggest a Poissonian process, validating approximations made in the model (see also fig. S4). **c.** MFPTs on the pillar array as a function of the pillar spacing (dotted). MFPTs of a Brownian particle with the same effective diffusion constant (*D* = 109 41*µm*^2^*/min*, fig 5**b**) as the cells (blue) exhibits the same trend as the measured data. When taking into account the cell-pillar interactions, by substracting the contact time per MFPT, there is excellent agreement between the measured MFPTs (solid) and predicted MFPTs (blue). **d.** Contact-time is measured as the time spent by the cell center within a radius of 1*µm* around the pillar. **e.** Contact time of the cell-pillar interactions per MFPT for all three configurations.

At the spatial scale of a pillar domain, and time scale of an escaping cell, we consider that the dynamics of the cell center-of-mass is approximately Brownian. This is a consequence of both the inherent persistence length of cell trajectories on a flat surface, which is of similar length as a pillar domain (**S1**), and the additional randomness induced by cell-pillar interactions. Using a narrow escape model developed for random motion in a field of obstacles^43^, we estimate the MFPT 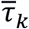 for a Brownian point particle escaping through funneled openings from a two-dimensional domain delimited by pillars as:

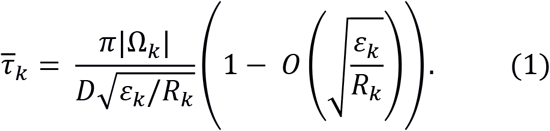

Here, *ε*_*k*_ is the space between pillars of radius *R*_*k*_ in the k^th^ domain (centered at *x*_*k*_) that delimit the area |Ω_*k*_|, and *D* is the diffusion constant of the escaping particle. In our assay, the pillar radius remains constant, so we express 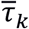 as a function of *L*_*k*_ = *ε*_*k*_ + *R*_*k*_, where *L*_*k*_ is the distance between two pillar centers (fig. **5a**). When the exits are sufficiently spaced, four similar escape routes are possible and thus the escape time changes to 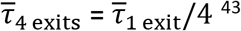^43^. The MFPT increases as pillars are spaced further apart, because both the pores widen (*ε*_*k*_) and the area of the domain (|Ω_*k*_|∼ *ε*_*k*_^2^) increases.

Figure **4c** shows the MFPTs (dotted line) measured for all cell trajectories with respect to the pillar spacing. The MFPT increases as pillars are spaced further apart. Furthermore, as *L*_*k*_ and |Ω_*k*_| are fully defined by the lattice, and the effective diffusion constant (*D*) is the only free parameter (eq. **1**), the expected MFPTs can thus be approximated using only the *D* found for starved *D. discoideum* moving on flat PDMS (**S1**). A linear fit to the mean-squared displacement (MSD) of the trajectories on flat PDMS allows to estimate the effective diffusion constant between the pillars is *D*_eff_ = 109 ± 41 µm^2^min^−1^ (**S1**, fig. **5b**). Then, the approximated MFPTs using this diffusion constant are shown in figure **4c** (blue line). Remarkably, the predicted and measured MFPTs share a similar slope when measured as a function of the inter-pillar spacing. As we will show in the next section, this slope can be linked to the topotactic drift, as it represents the difference in MFPTs across the spatial gradient.

**Figure 5:**
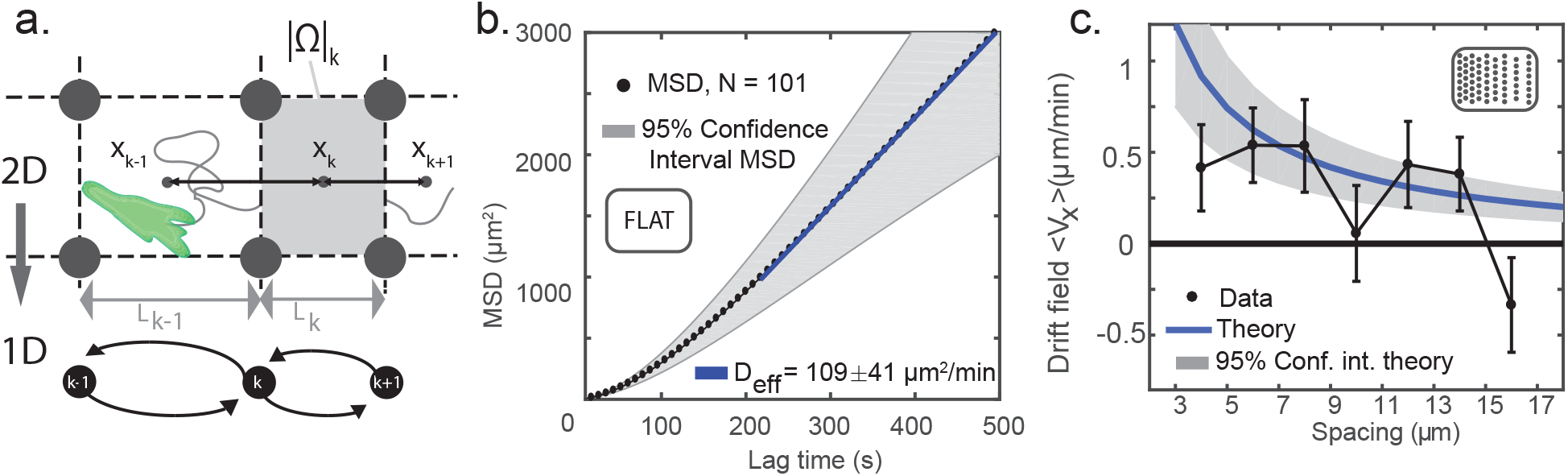
**a.** Schematic of modelling cell motion in a pillar array of asymmetric spacing. Cell centers are modelled by a diffusing particle escaping from domains (Ω_*k*_) delimeted by pillars (*r*_*k*_) of varying center-to-center distance (*L*_*k*_). To explain the drift we coarse grain this process to a one-dimensional non-isotropic random walk in the direction of the topotactic gradient. **b.** Approximation of the effective Diffusion constant (*D* = 109 ± 41*µm*^2^*/min*) by a linear fit to the MSD for lagtimes *>* 200 (see also fig. S1). **e.** The measured topotactic drift compared to the drift predicted by the model using the experimentally determined diffusion constant.

We note that there is a large offset between the measured and modelled MFPTs. Cells predominantly slide past pillars (fig. **1f** **Movie 1**), though sometimes, cells engage in long pillar interactions (**S5** and **Movie 5**). Such long interactions, although sporadic, can add a significant amount of time to the measured MFPTs, because cell transitions into new inter-pillar domains are delayed. The model assumes a reflective interaction and thus does not take into account these sporadic, but long interactions. To investigate if without these events the predicted and measured MFPTs agree, we measured this ‘dwell’ or contact time. We approximate the contact time as the time spent by a cell-center within a radius of 1 µm around a pillar (**S5**). Thereby, these contact times approximate the duration of long cell-pillar interactions, which are typified by cells primarily adhering to the wall of a pillar (fig. **4d**). The average contact times of the topotaxis experiment (T), and multi-cue experiments (O&A), are shown in figure **4e** and last 34.8 ± 2.6*s*, 54.8 ± 4.5*s* and 49.1 ± 4.7*s* per domain (Ω_*k*_) respectively. Interestingly, the multi-cue contact times are significantly longer. This can be attributed to the overall lower speed of *D. discoideum* when exposed to cAMP gradients (**S1** and ^44^). When these average contact times are subtracted from the measured MFPTs, the resultant MFPTs give a very good fit to the theory (fig. **4c**, solid line).

### Stochasticity of cell motion results in topographical guidance on an asymmetric pillar lattice

So far, we showed that amoeboid migration in an asymmetric pillar field results in topographical guidance and that this effect is conserved during chemotaxis. Furthermore, as the pillar field induces enough randomness in the cell trajectories, the MFPTs can be approximated by Brownian trajectories with a similar diffusion constant. We then showed that the MFPTs of the model and experimental data share the same slope when plotted as a function of the pillar spacing. Building on this crucial observation, we now propose that topographic guidance in the pillar field is a result of different escape rates (or MFPTs) across domains of inter-pillar spacing. To show this, we calculated the net flux of Brownian trajectories in the pillar lattice with a spatial gradient, and then use this calculation, to compare the strength and direction of the modelled and measured topotactic drift of amoeboid cell migration.

We coarse-grain the dynamics of cell migration in the pillar lattice as a random walk between centers of adjacent domains in the direction of the gradient (fig. **5a**). In this model the cell jumps to the next site at exponentially distributed waiting times, with different escape rates. The previously introduced MFPT (eq. **1**) gives the escape rate towards the more (*k* − 1) or less spacious (*k* + 1) side:

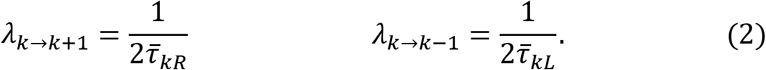

Using these escape rates as a basis for the transition probabilities, we then approximate the master equation for the transition probability density function of the random walk by a two-dimensional convection-diffusion equation and derive a one-dimensional equation by projection onto the x-axis (**S6**). From the master equation we obtain the following Fokker-Planck equation on the lattice:

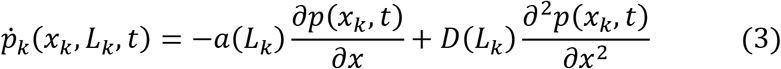

Here, *x*_*k*_ is a spatial coordinate, with *x*_*k*+1_ = *x*_*k*_ + *L*_*k*_ and *x*_*k*−1_ = *x*_*k*_ − *L*_*k*_, where again, *L*_*k*_ is the pillar center-to-center distance. The effective drift *a*(*L*_*k*_) and effective diffusion constant *D*(*L*_*k*_), can be used for a coarse-grained description of the random walk between domains:

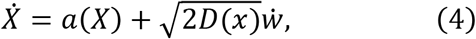

where 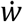 is the Wiener process. Hence, the difference in escape times towards and away from the more spacious and less spacious sides (**S6**), results in an effective topotactic drift *a*(*L*_*k*_) for a random walker between domains,

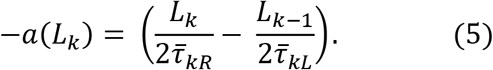

In deriving the model above, two key approximations were made. First, the MFPT from each domain must follow an exponential distribution in order to model the movement as a stochastic process on the lattice. The escape time measurements, shown in figure **4b** (and **S4**), confirm that this is indeed the case. Secondly, the MFPT of the cell center, escaping from a domain, should on average be approximated by a diffusing particle with diffusion constant *D* (eq. **1**). In the previous section, we showed that although the measured MFPTs are much longer than predicted (when using *D*_eff,flat_), the slope of the MFPTs as a function of pillar spacing is similar. Since only the difference between MFPTs enters equation (5), the drift predicted by our model depends exclusively on this slope. Hence, the topotactic drift is unaffected when adding a constant contact time.

To predict topotactic drift (eq. **5**) we needed to estimate the diffusion constant *D* of the cells on a flat surface, as the other variables determining the MFPT (eq. **1**), the domain size |Ω_*k*_|, and the pillar spacing *L*_*k*_, are determined by the geometry of the lattice. As mentioned before, we approximate the diffusion constant by a linear fit on the average MSDs found on flat PDMS (fig. **5b** and **S1**). Figure **5b** shows the MSDs of 101 cell trajectories and their average (black) as a function of the lag time *τ*. On short time scales (< 80 s) cell movement consists of random and persistent motion (**S1**, *a* ≈ 1.7), at intermediate lag times (80-200 s) randomness starts to dominate, resulting in purely diffusive behavior at long time scales (>200 s). The measured MFPTs are of the same order of magnitude as the intermediate to long lag time regimes of the MSD.

Using the lattice constants (|Ω_*k*_|, *ε*_*k*_, *R*_*k*_) and the approximation of the diffusion constant (*D*_*eff*_), we compare our predictions (eq. **4**) of the drift on the lattice to the experimental data. Figure **5c** shows that the predicted drift (blue line) adequately describes the measured topotactic drift for most pillar spacings (5 − 15*μm*). This suggests that the topotactic drift originates from the combination of the pillar lattice with the inherent, random component of cell motion. We note that chemotaxis has a large random component (see fig. **S1-2**), thus the topotactic drift should equally affect chemotaxing cells.

## Discussion and conclusion

We investigated highly motile amoeboid cell migration in a multi-cue environment, using the model organism *D. discoideum* in the starved state. An anisotropic pillar field with a gradient in inter-pillar spacing was demonstrated to generate a topographical cue, while preserving the characteristic motion of the migrating cells. With an added chemical gradient, the cells were exposed to two simultaneous external stimuli. By performing experiments with the chemical and topographical gradients either aligned or opposed, we found that the directional bias due to the topographic asymmetry is conserved when the cells undergo chemotaxis. The results of these experiments show that the topotactic and chemotactic effects add up linearly.

In the second part of this paper, we studied the origin of the topotactic effect by modelling cells in the pillar field as diffusing particles escaping from pillar domains. The differences between MFPTs of such particles, towards and away from wider pores and more spacious areas, were shown to result in topographical guidance. Precisely these differences in modelled MFPTs agree with the experimental measurements, enabling an adequate prediction of both the strength and direction of this particular long-range topotactic drift. This form of topotaxis is conserved in multi-cue environments as it is an emergent property of the combination of asymmetric spatial constraints of the lattice and the inherent stochasticity of (biased) amoeboid migration. Surprisingly, the differences in MFPTs between pillar spacings can be modelled using Brownian dynamics, since the cells clearly have a persistent component to their motion when moving on a flat surface. The interplay between the topography and the cell must induce enough randomness, thus lengthening quasi-random motion, to allow for such a model. Considering the cells do retain some persistence, it would be interesting to investigate the contribution of persistence to topotaxis generated by asymmetric lattices. Finally, the success of a diffusion model suggests that the drift induced by the topography is a process with no memory.

Until now, few *in vitro* experiments have been performed with two well-controlled simultaneous stimuli. For the combination of topographical and chemical guidance specifically, we are only aware of the work by *Comelles et. al.*^17^ (hapto-vs topotaxis). Although not explicitly analyzed as such in their study, the presented cell migration data in an assay consisting of both a ratchet (topotaxis) and a fibronectin gradient (adhesional haptotaxis) seems to indicate linear addition of these cues as well. It is important to note that their assay also does not confine cell motion to one-dimension, unlike many other forms of topographical guidance studied so far^15,16,45^, and therefore allows for a natural, unconstrained haptotactic response. Building on the results presented here and by *Comelles et. al.* a next step could be to investigate other topographic effects combined with chemical gradients, starting with the smaller than cell size topographies in which *Park et al.* have recently reported topotactic effects to occur as well^22,23^.

To conclude, we showed that subtle topographic asymmetries can lead to a significant influence on the chemotactic response of cells. This observation is not only relevant for *in vitro* applications, like tissue engineering^46^ and cell sorting^47^, but also when studying amoeboid migration *in vivo*. Recent examples of such studies are on leukocyte migration^30^ and on the detachment of tumor cells^48^. While these *in vivo* studies usually focus on chemotaxis, our results suggest that an additional topotactic influence has to be considered as a key factor, and that it even might be necessary to correct for any of its effects. The results and model presented in this letter can thus serve as a stepping stone for future *in vivo* cell migration research.

## Materials and methods

### Cell culture and live cell imaging

The axenic *D. discoideum* (Ax2) with GFP insertion, strain HG1694, was obtained from Dr. Günther Gerisch (MPI for Biochemistry, Martinsried, Germany). The cells were grown at 20 °C in HL5 medium, supplemented with the antibiotic gentamicin at a concentration of 20 µg ml^−1^ (G-418, Biochrom AG, Berlin, Germany). Cells were cultured in 100 mm petri dishes (100mm TC-treated culture dish, Corning, Corning, USA) and confluency was kept below 70%. For microscopy experiments, cells were harvested and centrifuged at 1500 rpm for 3 minutes followed by three successive washing steps of the cellular pellet with 17 mM K-Na-phosphate buffered saline, adjusted to *pH* = 6.0 (PBS, Sigma Aldrich, Steinheim, Germany). After resuspension in PBS, the cells were placed in a conical tube in a shaker at 20°C for 30 minutes. To induce cAR1 expression, cells were then pulsed with 150 nM cAMP applied in 6 minute intervals over 4 hours while shaking. Any residual cAMP was then removed after centrifugation and after resuspension the cells were left to shake in a conical tube with PBS for another 30 minutes.

Post-pulsing, cells were applied to the microfluidic observation chamber in PBS solution. The cell suspension was added to the chamber (sticky Slide I 0.8 Luer: sterilized, Ibidi, Gräfelfing, Germany) until a concentration of less than 10 cells per 360 by 360 µm (camera field of view) was found to adhere to the surface. Imaging started 1 hour after seeding the cells in the chamber. In the case of a combined gradient, cAMP was pumped (Tubing: BIO-rad, Hercules, USA. Syringe: 710 LT, Hamilton, Bonaduz, Switzerland) into the microfluidic channel of the observation chamber during this time and let to diffuse (**S6**), establishing a chemical gradient.

Cells were imaged every 8-10 seconds for 1-2 hours on a Nikon Eclipse Ti microscope equipped with a Yokogawa confocal spinning disk unit operated at 10,000 rpm (Nikon, Japan). GFP was excited by a 488 solid state diode laser (Coherent, Santa Clara, CA, U.S.A.), supported in an Agilent MLC4 unit (Agilent Technologies, Santa Clara, CA, U.S.A.). Images were captured by an Andor iXon Ultra 897 High Speed EM-CCD camera (Andor Technology, Belfast, Northern Ireland).

### Fabrication of pillar gradient structures

PDMS (Sylgard 184 Silicon Elastomer Kit, Dow Corning, MI, USA) micro-pillars were made as previously described in ^41^ using standard photolithography techniques. The micro-pillars were positioned in a trigonal array. The center-to-center distance was varied every four pillars in one direction such that each array has regions of low to high density. Pillar centers range from 11-25 µm. The pillar diameters were kept constant at 8 µm, resulting in an effective change in inter-pillar distance (pillar wall to wall) of 3-17 µm. The PDMS was hydrophilicitized by 10 min of UV/Ozone exposure (UVO-42, Jelight Company, Irvine, CA, U.S.A). Subsequently, the micropillar arrays were washed 3 times with Ethanol (70%) and PBS respectively. The PDMS pillar field was placed at a distance of 20 ± 0.5 mm from the inlet of the microfluidic channel.

### Establishing the chemotactic gradient

The PDMS arrays were arranged in parallel or in an anti-parallel direction with respect to the established cAMP gradient. To achieve an optimal chemotactic response from the cells, the cAMP gradient (∇*c*) was kept between 10^−3^ − 10 nM during experimentation. The gradient was set up by injection and subsequent diffusion of 40 µL of 10^−4^ M cAMP in PBS inside the microfluidic observation. The precise volume of cAMP solution needed was calibrated using a fluorescent dye (Alexa fluor 488 Hydrazide, Thermo-Fisher, Waltham, USA) which has a similar molecular weight to cAMP, 570.48 to 329.20 g/mol respectively (fig. **S8**). Additionally, finite element simulations were performed with COMSOL MultiPhysics (COMSOL BV, Zoetermeer, NL) to investigate the stability of the gradient over time, to determine suitable injection concentrations and to explore any effects of the pillar geometries on setting up a gradient by diffusion (fig. **S9**)

### Cell tracking

Cell movement was analyzed by rendering each image in the time-series into a binary image in ImageJ (http://imagej.nih.gov/ij/) and using the CellEvaluator plugin^49^ to determine center-of-mass xy-coordinates of each cell object per frame. We analyzed the xy-trajectories with in-house Matlab (version 2018b, The Mathworks, Inc., Natick, MA, U.S.A.) analysis program. Using the recorded displacements, and registered imaging times, we calculated movement statistics, like instantaneous velocities, turning angles and mean-square displacements (MSD), as previously published^38,41^.

To prevent a bias in the analysis by still unstarved, dead or otherwise immotile cells in the data set, an MSD selection criterion was employed. All cell trajectories that had an MSD (∆*R*^2^) after a lag-time of *τ*_*crit*_ = 200 *s* of

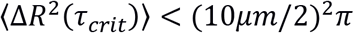

or lower, were deemed too passive to be of interest for this motility study. Note that a typical cell has a diameter of 8 − 12*μm* ^40^.

### Cell motion as a function of topography

To analyze the behavior of the cells as a function of topography, the pillar field was either dissected in bands of equal spacing (fig. **3b.**) or divided into regions based on the pillar lattice (fig. 4**a.**). To identify regions of interest, the position of each pillar was identified using pillar images recorded immediately before and after each experiment. After processing the images (binarizing and applying a Gaussian blur: *σ* = 2), pillars were recognized using appropriate Sobel filters using a custom MATLAB algorithm.

For the transition probabilities between different pillar spacings (fig. **3d-f.**), all transitions of cells from band-to-band were counted (*T*_*n*_). The bands were separated by linking all pillar-centers (in the y-direction) by a linear fit, generating pillar lines in y. Subsequently, a transition was defined as the cell center crossing past a pillar line by 2 µm. Transitions were then separated between going towards the more spacious (+*x*, *T*_n,spacious_) or less spacious (−*x*, *T*_n,narrow_) part of the array. The transition probabilities were then calculated by:

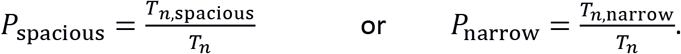

To calculate the drift as a function of spacing (fig **2e-g**), all instantaneous velocities (*ν*_*x*,*i*_) in the direction of the gradient were summed over for that pillar spacing: 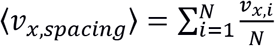.

For the MFPT analysis (fig. **4a-b.**), the quadrilateral lattice was constructed by linking all pillar centers in four directions, using a nearest neighbor algorithm, dividing the pillar field in domains delimited by pillars. A transition was recorded if the cell center crossed the division between domains by 1 µm. The directions as well as the time between escapes were recorded. Because the pillar lattice changes spacing, lattice defects can form, sometimes leading to irregularly shaped small and large domains, escapes from such regions were discarded from the analysis. To estimate the time cells interact or ‘stick’ to pillars, a dwell time was calculated. Using the positions of pillar centers obtained, the time that a cell center continuously dwells in a radius of maximally 1 µm outside of each pillar was recorded.

### Determining average values and their error estimates

We grouped measurements of observables over similar interpillar distances. This way probability density functions were constructed of observables at certain spacing, like the drift (fig. **2e-g**) and escape time (fig. **4c**). Such distributions were characterized by their average 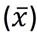. Error estimates were based on 95% confidence intervals using a t-distribution,

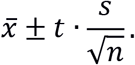

Here *t* is the upper critical value for the t-distribution (*t* = *F*^−1^(*p*|*ν*)), where we use *p* = 0.025/0.0975 and *ν* = *n* − 1, where *ν* are the degrees of freedom and *n* is the sample size. The two-state duration and escape time probability density functions were fit to exponential distributions using the MATLAB curve fitting toolbox with 95% confidence intervals on the fit parameters. In these fits, the fitting parameter *τ* was defined as *f*(*t*) = *AA*^−*λt*^, with 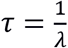.

