## Supplementary material for "Chemotaxis and topotaxis add vectorially for amoeboid cell migration"

#### Contents

**1 Motion of starved *D. discoideum* on flat PDMS surfaces 2**

**2 Comparison of the average drifts for all configurations 6**

**3 Cell motion in a steep topotactic gradient 7**

**4 Escape times of migrating cells in a trigonal pillar lattice 8**

**5 Contact time of long cell-pillar interactions 9**

**6 Modelling topotaxis in a pillar lattice 10**

**7 cAMP gradient formation in a microfluidic containing PDMS micropillars 12**

### 1 Motion of starved *D. discoideum* on flat PDMS surfaces

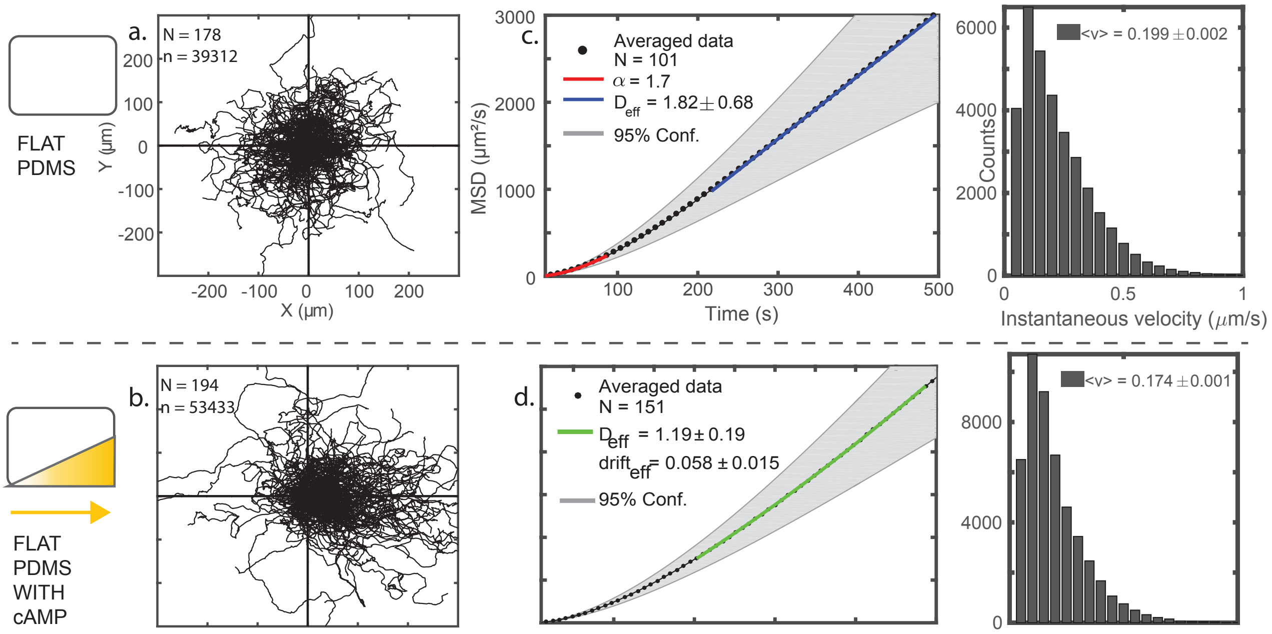


Figure S1: **Cell migration of starved *D. discoideum* on flat PDMS with and without a cAMP gradient. a.** Cell trajectories measured on flat PDMS without a cAMP gradient. **b.** Cell trajectories measured on flat PDMS with a cAMP gradient (source at $+x$). **c.** The mean-squared displacement of cell trajectories on flat PDMS. A power-law fit as a function of time ($\leq$ 80 seconds) is shown in red and a linear fit to extract the diffusion constant ($>$ 200 seconds) is shown in blue. **d.** The mean-squared displacement of the cell trajectories on flat PDMS with a cAMP gradient. A fit to the biased random walk model is shown in green. **e.** Histogram of the instantanous velocity measured for all trajectories on flat PDMS. **f.** Histogram of the instantanous velocity measured for all cell migration trajectories on flat PDMS in the presence of a cAMP gradient.

To benchmark the investigation of cell movement through an anisotropic pillar lattice and a chemical gradient, the motility of *D. discoideum* was first characterized on flat surfaces of Polydimethylsiloxane (PDMS). This is important, as although *Dictyostelium discoideum* (*D. discoideum*) migration has been characterized on flat PDMS for vegetative cells [1], when the cells are starved, and become sensitive to cyclic-adenosine monophospohate (cAMP) chemotaxis, their movement changes drastically. Previous measurements on different materials have shown that both migratory speed and persistence increase significantly [2].

#### 1.1 MSDs of cell trajetories with and without a chemical cue

Recent theories modeling *D. discoideum* motility have mostly focused on the vegetative state of the amoeba [8]. It is important to note that in the starved state cells move markedly different. Like previous studies [2], we also find that both speed and persistence increase of starved *D. discoideum*. When in this starved state, the cells try to pick up on cAMP cues to form aggregates. However, when devoid of such a cue, they walk around randomly to pick up on a signal, and while doing so, the statistics of their trajectories resemble those of persistent random walkers (PRW). The MSD of a PRW is $\langle r(t)^2\rangle = 2v_c^2\tau_p^2(t/\tau_p + e^{-t/\tau_p} - 1)$, where $v_c$ is a constant velocity and $\tau_p$ is the persistence time of the walker. This expression for the MSD has limiting behavior for short times ($t \ll \tau_p$) of $\langle r(t)^2\rangle = (v_c t)^2$ and long times ($t \gg \tau_p$) of $\langle r(t)^2\rangle = 2v_c^2\tau_p t$. In other words, it describes purely ballistic motion at short times and purely diffusive behavior at large times.

The average mean-squared displacement (MSD) of 178 trajectories measured on flat PDMS (figure S1 **c.**) resembles a curve that would be obtained for persistent random walkers [6]. However, a power-law fit ($\langle r(t)^2 \rangle = At^{\alpha}$) reveals that on short time scales (< 100s) the motion is not purely ballistic ($\alpha \approx 1.7$ versus $\alpha = 2$). The ballistic-like regime is followed by an intermediate region (100 − 200s, for which $\alpha \neq 1$) until the purely diffusive regime is reached (> 200s). A linear fit of the diffusive regime returns an effective diffusion constant of $D_{eff} = 109 \pm 40.8$ $\mu m^2 min^{-1}$ ($1.82 \pm 0.68$ $\mu m^2 s^{-1}$). For long times, the measured MSDs grow linearly with time, congruent to a PRW. Concluding, although qualitatively the MSDs indicate persistent random motion, the cell centers do not perfectly follow a PRW. This observation parallels results found in other experiments and for other cell types [9][10][11].

Then, to compare cell response in a multi-cue environment to only chemotaxis, 194 cell trajectories were recorded in a cAMP gradient on a flat surface (figure S1**b**). The cell trajectories show that the cAMP gradient in our setup (MM and S7) has a clear chemotactic response and lies in the medium signal-to-noise regime [5] of what is maximally achievable for *D. discoideum*. The MSDs of chemotaxing cells show a similar ballistic-like regime at short timescales, evolving in a diffusive regime at long times. However, for chemotaxing cells there is an overlaid drift ($v_d$), the MSD curve is not linear for long times, and thus the data seems to indicate a biased random walk (BRW). The MSD of a BRW is given by: $\langle r(t)^2 \rangle = (v_d t)^2 + 4Dt$. A fit to this model returns an effective diffusion constant of $D_{eff} = 71.4 \pm 11.4$ $\mu m^2 min^{-1}$ ($1.19 \pm 0.19$ $\mu m^2 s^{-1}$) and drift velocity of $v_d = 3.48 \pm 0.90$ $\mu m min^{-1}$ ($0.058 \pm 0.015$ $\mu m s^{-1}$). All error margins to the MSD fits are defined by additionally fitting to the minimum and maximum of 95% confidence bounds of the averaged MSD (grey, fig. S1**a-b**). Finally, the instantaneous velocities of starved *D. discoideum* moving on flat PDMS measure $\langle v_{inst} \rangle = 11.9 \pm 0.1$ $\mu m min^{-1}$ ($0.19 \pm 0.002$ $\mu m s^{-1}$) and $\langle v_{inst} \rangle = 10.4 \pm 0.1$ $\mu m min^{-1}$ ($0.17 \pm 0.001 \mu$ $m s^{-1}$) without and with a cAMP gradient respectively. The drop in speed for cells exposed to a cAMP gradient has been previously reported [7].

Although the cells move in a persistent-random way, their motion cannot be fully described by, or fit to, the PRW model. However, it is instructive to estimate the trajectories average persistence. The measured MSD of starved cells without (fig S1**c**) and with (fig S1**d**) a chemical cue develop into a linear function between $100 - 200s$. Combining this observation with the average instantaneous velocities in both experiments, the persistence length of trajectories average between $20 - 40$ $\mu m$ and $17 - 34$ $\mu m$ respectively. This means the persistence of cell center trajectories is around two to four cell lengths.

#### 1.2 Local-MSD analysis of cell trajectories with and without a chemical cue

The previous section analyzed starved *D. discoideum* trajectories by comparing various motility models to global MSDs. For long lag-times, movement is diffusive on flat PDMS, and drift-diffusive after adding a chemical gradient. However, on short to medium time scales (< 200s) it is unclear what contributes to the non-linear part of the MSD curve. At short times, multiple processes might be at play, each dominating at different time scales. The local-MSD (l-MSD) analysis is an alternative technique to study persistence during cell migration, and well suited to study cross-overs between ballistic and random motion. This analysis can be used to distinguish between directed and quasi-random motion states within one trajectory. For vegetative *D. discoideum* cells, this method was previously published in [1].

For every time point $t_i$ in the trajectory, representing the center of a rolling time window ($T = M\delta t, M = 12$), we first calculate and then fit the l-MSD to a local lag-time:

$$\Delta R^2(t_i, \tau_k) = A\dot{\tau}_k^{(\alpha_i)} \quad (1)$$

The exponent $\alpha$ in equation 1.2 characterizes the migratory state of the cell at time $t_i$. The prefactor A is either a diffusion coefficient analogue, when in a quasi-random motion state, or can be interpreted as the velocity in the case of directed motion. Additionally, we calculate the angle persistence function $\Delta\phi_i$ (as defined in [1]) inside the same rolling window.

A time point is assigned to a directed-state when $\alpha \sim 2$ and the value of the angle persistence function stays within pre-defined bounds:

$$P_{i,dir} = \begin{cases} 1 & \text{if } [2 - \sigma_\alpha \leq \alpha_i \leq 2] \cap [0 \leq \Delta\phi_i \leq \sigma_\phi] \\ 0 & \text{otherwise} \end{cases} \tag{2}$$

To allow for the zig-zag motion that cells exhibit on short time-scales when directionally travelling [8], we choose $\sigma_\phi = 3\sigma_\alpha$ and $\sigma_\alpha = 0.3$.

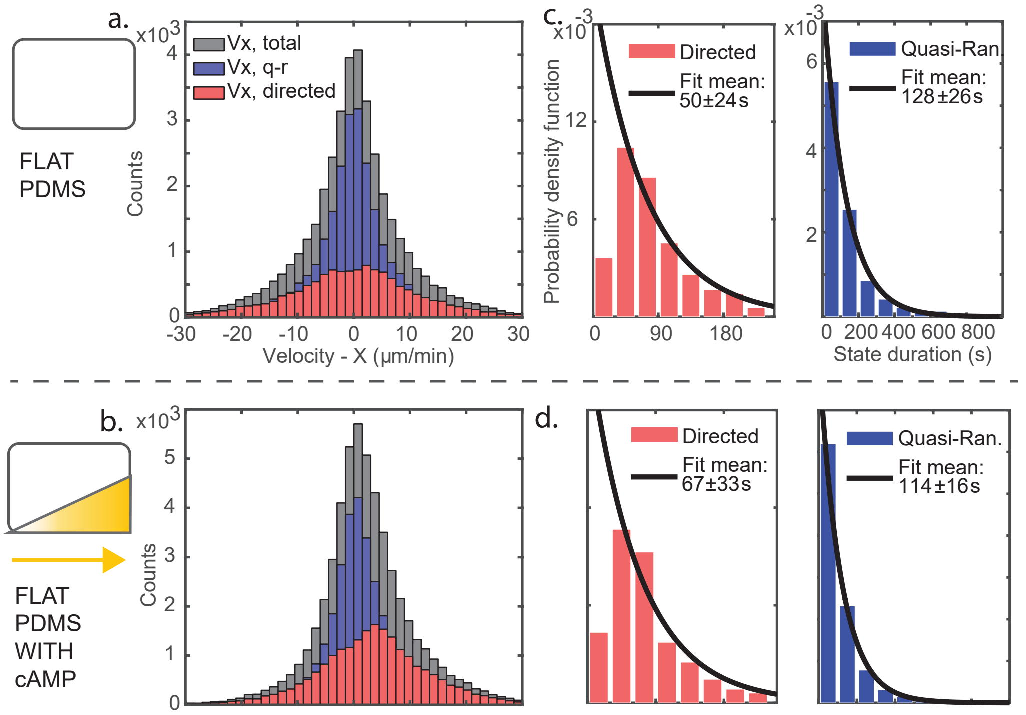


Figure S2: **l-MSD analysis of cell trajectories on flat PDMS with and without cAMP.** **a.** The X-component of the velocity for cells moving on flat PDMS (grey), without a chemical gradient. Measured velocities are also split for directed runs (red) quasi-random motion states (blue). **b.** The X-component of the velocity for cells moving on flat PDMS (grey), with a chemical gradient. **c.** State durations of starved *D. discoideum* in either the directed run (red) or a quasi-random motion state (blue) on flat PDMS. The probability distributions are fit to an exponential decay. **d.** Measurement of the state durations of cells subjected to a cAMP gradient.

Using this technique, we first study the drift velocity ($v_x$). On a flat surface the starved cells have a negligible drift in the $x$-direction (direction of cues). The velocity distributions (figure S2**a**) were fit to a Gaussian for all data ($\mu = 0.038 \pm 0.35$ $\mu mmin^{-1}$, $\sigma = 8.2 \pm 0.6$), for directed motion states ($\mu = 0.11 \pm 0.18$ $\mu mmin^{-1}$, $\sigma = 14.6 \pm 0.6$) and quasi-random motion states ($\mu = 0.01 \pm 0.31$ $\mu mmin^{-1}$, $\sigma = 6.1 \pm 0.4$). The distributions show that there is a negligible drift for these trajectories and this observation is conserved when separated in directed and quasi-random motion states. The opposite is measured for cells subjected to a cAMP gradient; the velocity distributions (figure S2**b**) are asymmetric with means (95% C.I., see MM) for all data: $\mu = 2.3 \pm 0.1$ $\mu mmin^{-1}$, directed runs only: $\mu = 3.6 \pm 0.1$ $\mu mmin^{-1}$ and quasi-random motion: $\mu = 1.3 \pm 0.1$ $\mu mmin^{-1}$. These means indicate a drift in the positive $x$-direction (direction of cAMP source). There is a difference between the average drift ($\langle v_x \rangle$, fig S2) and the one found by the BRW fit. Two things contribute to this difference. First, the BRW fit performed does not include for short lag-time persistence. Secondly, there is a difference in both data sets, as trajectories that are more biased leave our field of view more quickly and thus do not have enough length to be included in a MSD analysis ($> 100$ datapoints per trajectory).

The probability distributions of the state durations of the cells on flat PDMS without and with a chemical gradient are shown in figure S2**c** and **d** respectively. The distributions are well fit by an exponential function ($f(t) = A\exp^{-t/\beta}$) giving means $\beta = 50 \pm 24s$ and $\beta = 128 \pm 26s$ for the directed and quasi-random states on a flat surface, $\beta = 67 \pm 33s$ and $\beta = 114 \pm 16s$ for the trajectories under influence of a chemical gradient. The shape of these distributions suggest a Poissonian process for the switching between states. The average duration of the directed state increases when the cells undergo chemotaxis, but not significantly. This suggests that it is not predominantly the change in persistence that generates the chemotactic drift, unlike the recently suggested mechanism for durotaxis [12], but rather a bias in reorientation of the cells.

Interestingly, the separation in 'directed' and 'quasi-random' motion states reveals that drift is highest in those parts of the trajectories that were designated as a 'directed state'. This seems intuitive, as these states select for directed movement. However, such a state should consistently point along the gradient direction to generate large drift. Combined with the observation that chemotaxing cells do not have significantly longer directed states (fig S2**c** and **d**), then suggests that the origin of chemotactic drift is reorientation in the quasi-random motion state. In varied signal-to-noise environments, generated by our setup, chemotaxis is a process primarily based on directional re-orientation, rather than the lengthening of directed runs. Finally, for estimating the average persistence length of trajectories, local-MSD analysis agrees with results of supplementary section 1.1. The duration of a cycle including one directed and one quasi-random state is between $100 - 200s$ on average, multiplied with the instantaneous velocities (fig S1**e** and **f**) we again retrieve a persistence length of around two to four cell lengths.

#### 2 Comparison of the average drifts for all configurations

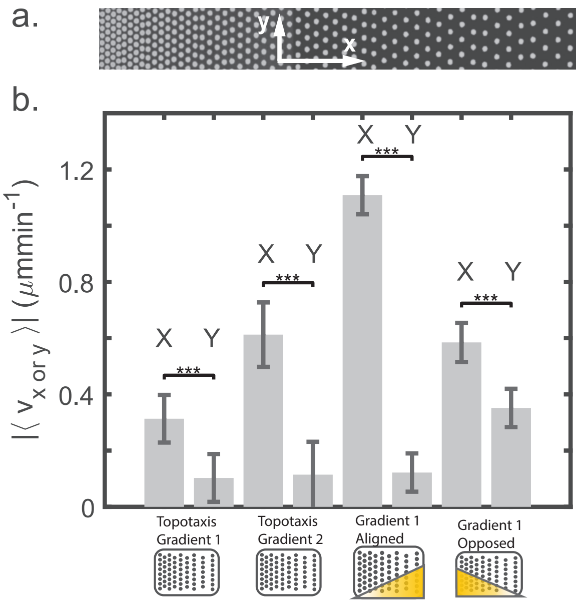


Figure S3: **Average drifts in direction of gradients are significantly higher than noise** **a.** The orientation of the directions $x$,$y$ with respect to the pillar array (topotactic gradient 1). **b.** The absolute value of the average drifts $|\langle v_{\mathrm{x\ or\ y}}\rangle|$ measured for all configurations in $x$ and $y$. The drifts in the direction of the gradients ($\pm x$) is significantly higher than the 'random' drift measured in the perpendicular direction ($\pm y$). Significance tests performed with a students $t$-test.

Figure S3 compares all the average drifts measured in directions $\pm x, \pm y$. The drifts in the direction of the gradients ($\pm x$) are significantly higher than in the perpendicular directions ($\pm y$). The average topotaxis in the first, more shallow gradient, is lower than in the steeper gradient (2). This is also obvious from the more detailed drift analysis (as a function of pillar spacing) shown in figures S4 and 2, where we can see that the drift in the shallow gradient has a turning point at spacing $15 - 17\mu m$ and the steep gradient does not. The opposed configuration has a notably higher drift in the $y$ direction than the other configurations. We speculate that this may be due to the cells making more use of straight alleys between pillars in $\pm y$, amplifying any random drift, which could be a consequence of a configuration with two conflicting gradients. This effect could be the topic of another study.

### 3 Cell motion in a steep topotactic gradient

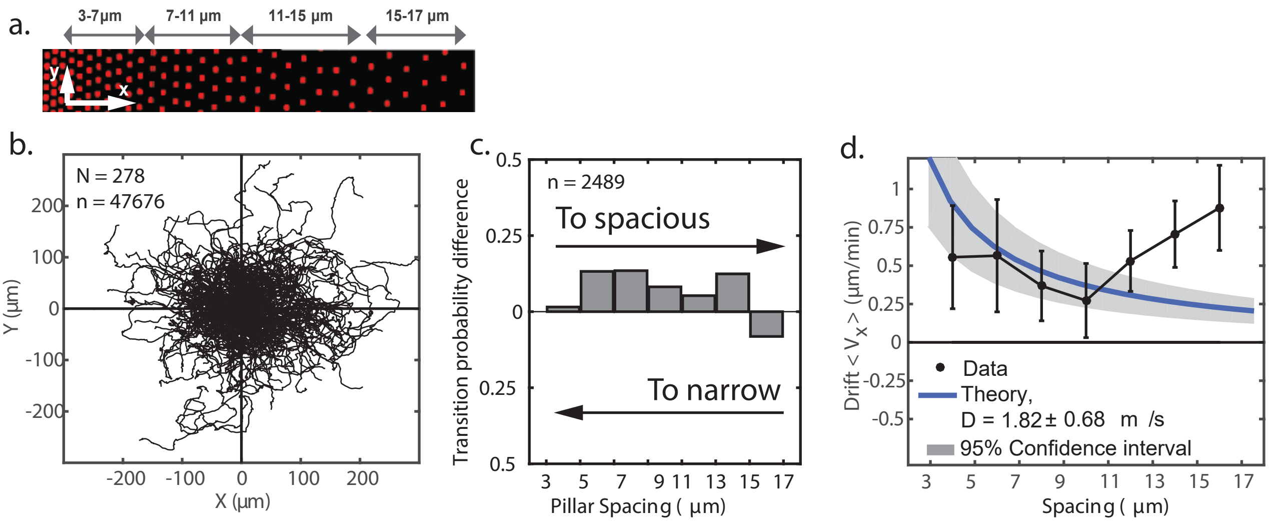


Figure S4: **Topotaxis on a steep spatial gradient**
**a.** Image of the *steep gradient* pillar array, tops of pillars labelled with DiI (RFP). **b.** Trajectories measured on this pillar field. **c.** Transition probabilities as a function of the spacing between pillars. **d.** The averaged drift field of the cells as a function of the spacing between pillars. The theoretical prediction based on the model introduced in the text is shown in blue.

To investigate whether the topotactic effect is conserved for different spatial gradients, we measured cell motility on an array with a steeper gradient. Figure S4**a** shows this array, where after each pillar row (in $y$) the pillar spacing changes by $1\mu m$ (in $x$). The 278 trajectories measured over 6 independent experiments on this pillar field are shown in figure S4**b**. The distribution of the trajectories is lob-sided towards the positive $x$-direction, which is the more spacious side of the array. This qualitative observation is confirmed by the transition probabilities found (figure S4 **c.**). Similar to the measurements performed on the array in the main text (figure 2**e**), on average, the cells tend to move towards the more spacious side of the pillar array, except for the largest spacings ($15 - 17\mu m$). The drift field as a function of the pillar spacing is shown in figure S4**d**. The error estimates (95% confidence) are significantly larger on this array than those measured on the other array (figure 2e). Additionally, the measured drift is only well described by the model (blue line) for small pillar spacings ($3 - 11\mu m$).

#### 4 Escape times of migrating cells in a trigonal pillar lattice

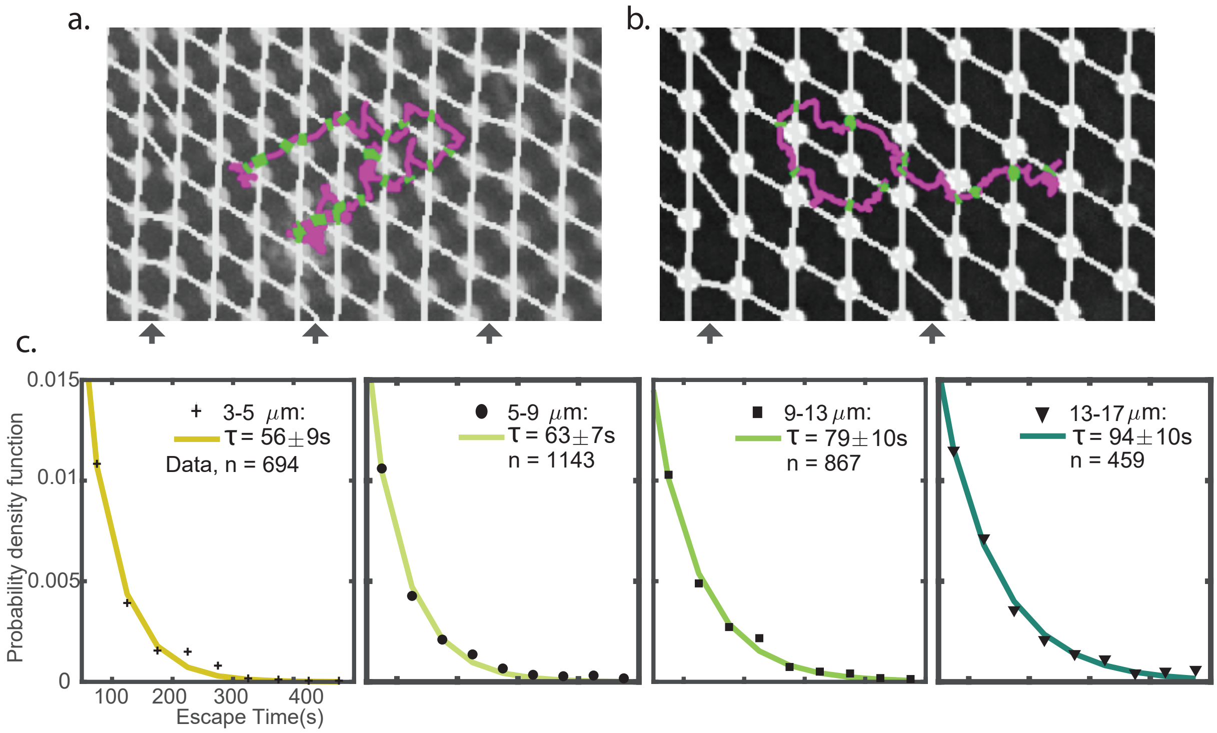


Figure S5: **Measured escape time distributions are well fit to an exponential, conserved across the entire field. a.** Trajectory of a cell-center (magenta) moving on a dense part of the pillar array. Each time the cell escapes a quadrilateral domain delimited by pillars the time is recorded (shown in green). Black arrows indicate where the spacing of the lattice changes. Scale bars represent $10\mu m$. **b.** An example of escape time measurements on a spacious part of the array. **c.** The escape time distributions are well fit by an exponentially decaying function for all spacing on the array ($2 - 17\mu m$). In each figure the mean of the fit with a 95% confidence interval ($\beta = 1/\lambda$) and number of data points (n) are specified.

Cell trajectories in the pillar lattice are effectively a series of escapes from domains delimited by pillars (figures S5 **a-b**). We measure the escape time by recording the time at which the cell enters a domain and then leaves again through one of four possible escape routes (figure 4**a**). These escape times can be very short, as a cell may turn and leave via the same entrance it came from, or very long as a cell touches pillars and explores the domain, leaving on the opposite side to where it entered.

When the lattice changes spacing (black arrows, fig. S5 **a-b** ), lattice defects arise in the connective network between the pillars. Such defects result in domains with three or five pillars along the border of a corresponding unit cell, giving extraordinarily large or small domain areas. All data involving such irregular unit cells are excluded from the mean first passage times (MFPT) data.

The MFPT measured on each spacing of the pillar field are well fit to an exponential distribution (figures S5 **a-b**). This suggests that the cell escaping from each domain is a Poissonian process. Interestingly, although a cell can have directed motion states larger than the typical size of a domain, on average the escape times measured suggest that the escape is a random process.

### 5 Contact time of long cell-pillar interactions

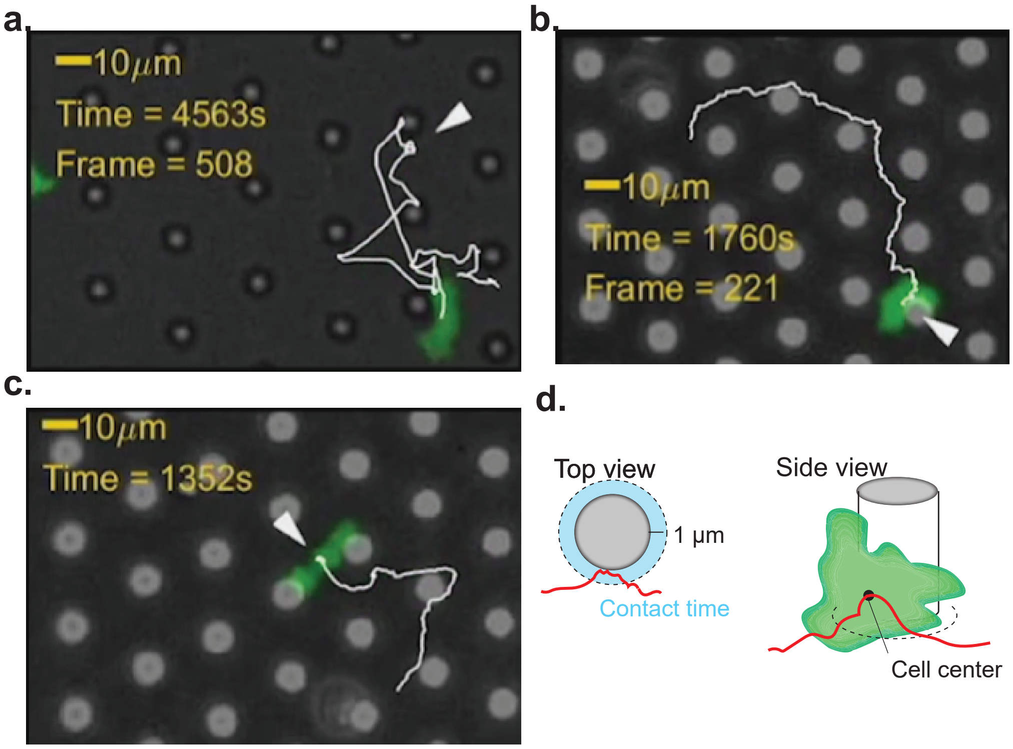


Figure S6: **Examples of long cell-pillar interactions. a. - b.** Cells predominantly slide past or bounce from pillars (shown in the figures by smooth trajectories), but sporadically spend a longer time exploring vertical areas of pillars, here highlighted by white arrows (trajectories have many points inside a small area). **c.** Cell engaged in a 'split decision event' are in contact with multiple pillars and remain static for an extended perdiod of time (trajectories do not advance in the middle of a pillar unit cell). **d.** The 'dwell' or contact time is defined as the time between entry and exit of a cell trajectory in and out of 1 $\mu m$ radius around each pillar.

The cells predominantly brush past pillars, which leads to short-lived quasi-random motion states induced by the topography. The example trajectory in figure 1**f** (Movie 1) shows some of these short-lived states, a result of the cell sliding or bouncing off a pillar. Additionally, cells have sporadic, much longer interactions with pillars. For example, the cells occasionally explore the vertical pillar surfaces with the leading pseudopod (fig. S6**a**), fully wrap around pillars (fig. S6**b**) or cope with a 'split decision when in contact with two pillars at once (fig. S6**c**). These types of cell-pillar interactions are also shown in Movie 4. Vertical pillar interactions contribute to overall lower migration speeds in the lattice, when compared to migration on a flat surface [13].

The cell-pillar interactions, although only sporadically encountered, result in extended periods of 'contact time and significantly influence measured MFPTs. The contact time quantifies, on average, how much time *D. discoideum* interacts with a pillar and was defined as the time spent by the cell-center within a radius of $1\mu m$ around a pillar. In practice this means the cell is wrapped around a pillar or predominantly adhering to the vertical face of a pillar. Using this method of approximation, cells brushing past pillars are sometimes also counted as cell-pillar interactions and may, erroneously, contribute to the average contact time. This error is marginal, as we use a small radius around the pillar ($1\mu m$) and it is compensated for by not accounting for cells in a multi-pillar interaction, like those wedged between pillars (fig. S6**c**).

### 6 Modelling topotaxis in a pillar lattice

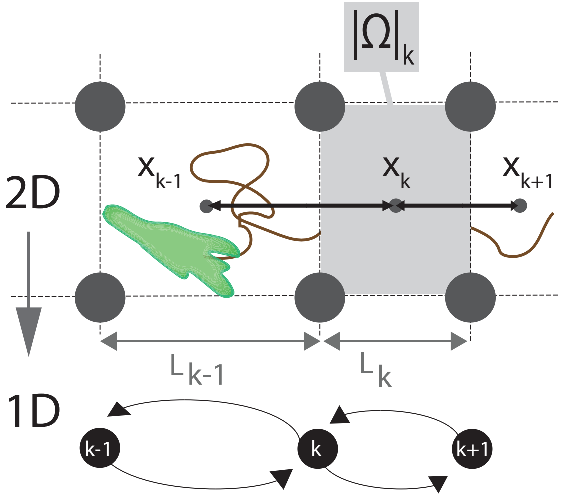


Figure S7: **Schematic depiction of a cell (green) moving through a pillar array with a spatial gradient** The center of each domain at spacing $k$ is denoted by $x_k$. The size of the domain is given by $|\Omega_k|$. The size of the opening between pillars is $\varepsilon_k = L_k - 2r_k$, where $L_k$ is the distance between pillar centers and $r_k$ the radius at spacing $k$ on the array.

#### 6.1 Drift of a stochastic process in a spatial density gradient

To characterize the dynamics of cells between pillars, we coarse-grained the cell motion as a jump process between the free available space delimited between pillars. The cell motion contains sufficient randomness, so that we model the motion using a stochastic equations. In particular, we shall compute the effective drift generated by the distribution of anisotropic pillar, that accumulate in the positive x-direction. There are no changes in pillar spacing in the y-direction and are uniformly distributed. The distance between neighboring pillars located at position $x_k$ is $dk$, their radii is $r_k$. The distance between two consecutive pillars is $L_k = |x_{k+1} - x_k|$.

#### 6.2 Derivation of a coarse-grained stochastic equation to model the random exploration of a cell inside a dense pillar field

We start with a continuous Brownian motion with diffusion coefficient $D$, the escape time between the funnel cusp-shape two pillars of size $R$ is

$$\bar{\tau} = \frac{\pi|\Omega|}{4D\sqrt{\epsilon/R}}\left(1 + O\left(\sqrt{\frac{\epsilon}{R}}\right)\right), \tag{3}$$

where the volume $|\Omega|$ and $\epsilon$ is the size of the opening [14]. The symmetry breaking between the pillar spacing on the right and on the left, lead to the two different exit times from the left and the right:

$$\bar{\tau}_{kL} = \frac{\pi|\Omega_k|}{2D\sqrt{\epsilon_k/R_k}}\left(1 + O\left(\sqrt{\frac{\epsilon_k}{R_k}}\right)\right), \tag{4}$$

and

$$\bar{\tau}_{kR} = \frac{\pi|\Omega_k|}{2D\sqrt{\epsilon_{k+1}/R_{k+1}}}\left(1 + O\left(\sqrt{\frac{\epsilon_{k+1}}{R_{k+1}}}\right)\right), \tag{5}$$

with $\epsilon_k = L_k - 2R_k$. Thus the rate of escape from left to right is the reciprocal of the mean time (the factor 2 accounts for the fact that half of the population can return)

$$\lambda_{k\to k+1} = \frac{1}{2\bar{\tau}_{kR}}, \tag{6}$$

and

$$\lambda_{k\to k-1} = \frac{1}{2\bar{\tau}_{kL}}. \tag{7}$$

The "Narrow Escape Theory" ensures that the escape distribution is Poissonian, it is then possible to compute the effective motion (diffusion coefficient and drift) of a coarse-grained stochastic process on a square lattice crowded by pillars.

We first coarse-grain the dynamics into a random walk between the centres of adjacent squares. Then we will approximate the master equation for the transition probability density function of the random walk by a two-dimensional convection-diffusion equation [15]. We will derive a one-dimensional equation by projecting on the $x$-axis. We recall that the mean exit time from a single lattice square is long, the first eigenvalue of the mixed Dirichlet-Neumann problem in a single cell is well separated from the higher ones. It follows that the waiting time is exponentially distributed with rates (6) and (7).

To derive the advection-diffusion equation from the master equation for the transition probability density function of a non-isotropic random walk that jumps at exponentially distributed waiting times on a square lattice with step size $L_k$.

The probability that the particle is in square $k$ (one dimension) at time $t+\Delta t$ is $p_k(t)$. The transition occurs from the square on the left with rate $\lambda_{k\to k-1}$ and probability $p_{k-1}(t-\Delta t)$, on the right with rate $\lambda_{k\to k+1}$ and probability $p_{k+1}(t-\Delta t)$ and finally the probability of no transition during $\Delta t$ is $1-\lambda_{k\to k-1}\Delta t + \lambda_{k\to k+1}\Delta t$ thus,

$$\dot{p}_k(t) = \frac{1}{2\bar{\tau}_{kL}} p_{k-1}(t) + \frac{1}{2\bar{\tau}_{k+1R}} p_{k+1}(t) - \left(\frac{1}{2\bar{\tau}_{kL}} + \frac{1}{2\bar{\tau}_{k+1R}}\right) p_k(t) \tag{8}$$

We shall now expand, in space, the density: $p_k(t) = p(x_k, t)$. Using a Taylor expansion with $x_{k+1} = x_k + L_k$ and $x_{k-1} = x_k - L_{k-1}$

$$p(x_{k+1}, t) = p(x_k, t) + L_k \frac{\partial p}{\partial x}(x_k, t) + \frac{1}{2} L_k^2 \frac{\partial^2 p}{\partial x^2}(x_k, t) + o(L_k^2) \tag{9}$$

$$p(x_{k-1}, t) = p(x_k, t) - L_{k-1} \frac{\partial p}{\partial x}(x_k, t) + \frac{1}{2} L_{k-1}^2 \frac{\partial^2 p}{\partial x^2}(x_k, t) + o(L_{k-1}^2) \tag{10}$$

Using a Markov chain [15], we obtain the coarse-grained Fokker-Planck equation on the lattice:

$$\dot{p}_k(t) = \left(\frac{L_k}{2\bar{\tau}_{kR}} - \frac{L_{k-1}}{2\bar{\tau}_{kL}}\right) \frac{\partial p}{\partial x}(x_k, t) + \frac{1}{2}\left(\frac{L_k^2}{2\bar{\tau}_{kR}} - \frac{L_{k-1}^2}{2\bar{\tau}_{kL}}\right) \frac{\partial^2 p}{\partial x^2}(x_k, t) \tag{11}$$

The effective drift $a(x)$ and diffusion tensor $D(x)$ are

$$a(x_k) = \left(\frac{L_k}{2\bar{\tau}_{kR}} - \frac{L_{k-1}}{2\bar{\tau}_{kL}}\right) \tag{12}$$

$$D(x_k) = \frac{1}{2}\left(\frac{L_k^2}{2\bar{\tau}_{kR}} - \frac{L_{k-1}^2}{2\bar{\tau}_{kL}}\right). \tag{13}$$

We use these quantity in the coarse-grained stochastic description which is valid at a spatial scale much larger than the space between two neighboring pillars:

$$\dot{X} = a(X) + \sqrt{2D(x)}\dot{w}, \tag{14}$$

where $w$ is the Wiener process.

### 7 cAMP gradient formation in a microfluidic containing PDMS micropillars

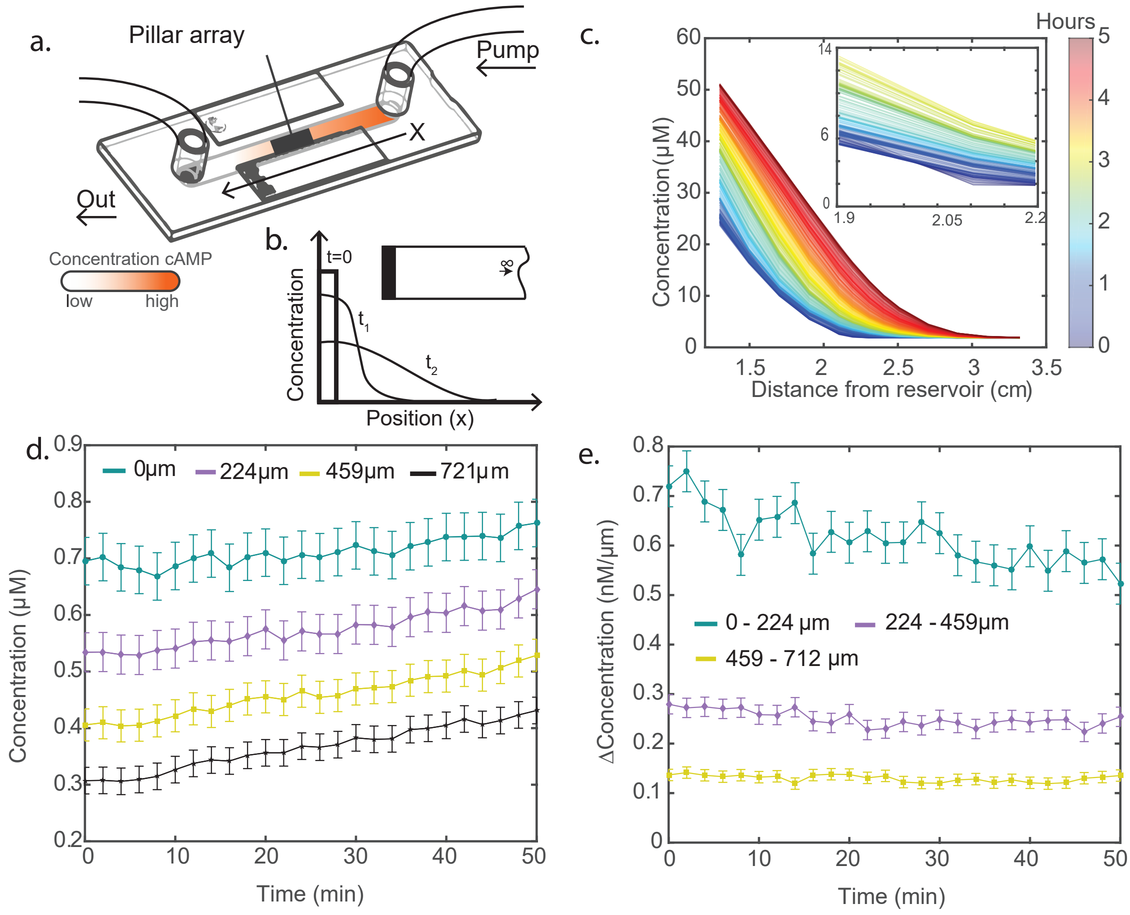


Figure S8: **Measurement of the cAMP gradient inside the microfluidic diffusion chamber**

**a.** Diagram of the microfluidic channel **b.** Qualitative sketch of the solutions to Fick's second law at different times for a solute diffusing into an empty semi-infinite bar (inset) **c.** Measurement of the diffusion of $75\mu L$ of the fluorescent dye Hydrazine 488 (similar molecular weight to cAMP) inside the microfluidic channel as a function of distance to the reservoir. **d.** Measurement of $40\mu L$ of the dye along the pillar field at different distances into the array, placed at 2cm ($0\mu m$ in graph) from the reservoir. Error bars are the standard deviation in measured fluorescence. During an experiment concentrations remain $\sim 0.1 - 1\mu M$ **e.** Difference in concentrations shown in **d.**, where the chemical gradient stays in the $\sim nM/\mu m$ range.

To establish the cyclic-adenosine monophosphate (cAMP) gradient over a PDMS micropillar array we used a microfluidic channel (figure S8**a**). The diffusion through the channel can be modeled by solving Fick's second law [16] in a semi-infinite bar (figure S8**b**):

$$\frac{\partial C(x,t)}{\partial t} = D\nabla^2 C(x,t) \qquad \int_0^\infty C(x,t)dx = B \qquad C(x,0) = 0$$

with the solution,

$$C(x,t) = \frac{B}{\sqrt{\pi D t}} \exp\frac{-x^2}{4Dt}.$$

Here $B$ is the total amount of solute, $D$ its diffusion coefficient, $x$ the distance from the initial boundary and $t$ the time. The chemical gradient is set up controllably, by slowly pumping in a set amount ($B$) of cAMP-PBS solution ($C$) and waiting a set time ($t$) for the solute to diffuse through the channel. By placing the micropillar array a particular distance from the reservoir ($x$), the chemical gradient can be replicated relatively well. However, the diffusion based microfluidic we used in this study does have less control over the chemical gradient than some more sophisticated, dynamic mixing, flow-based setups we [3] and others [4] have used in the past. A diffusion gradient was chosen over a mixing chamber, as a diffusive gradient is unaffected by obstacles (figure S9), where a mixing process is perturbed. A diffusion chamber also suffices, as *D. discoideum* exhibits chemotaxis in a very wide range of chemical gradients $\nabla C = 10^{-3} - 10\ nM$ (with a peak motility between $\nabla C = 0.01 - 0.1\ nM$) and over a similarly wide range of baseline concentrations ($\sim \mu M$) [4].

To benchmark the volume of solute, concentration, waiting time and distance from the reservoir, the diffusion of the fluorescent dye Hydrazine 488 was imaged inside the microfluidic channel (figure S8 **c.**). Hydrazine 488 has a similar molecular weight to cAMP, 329.206 g/mol versus 570.48 g/mol. The pillar array was placed at $x = 2$cm, before experimentation a volume of $B = 40\mu L$ with a cAMP concentration of $C = 10^{-4}M$ was pumped in slowly and imaging began after waiting $t = 60$min. During experimentation this yielded a cAMP gradient over the pillar field of $\sim 0.1\ nM$ (figure S8**d-e**).

To make sure the pillar array does not significantly influence the chemical gradient diffusing through the channel during the experiments, the setup was numerically investigated using finite-elements simulations (figure S9**a-c**). The chemical gradient is not significantly different at various heights inside and above the pillar field (figure S9**d**) and difference is equally negligble from the left to right (figure S9**f**) on the array. As expected, there is a gradient difference between the front and back of each pillar (figure S9**g**).

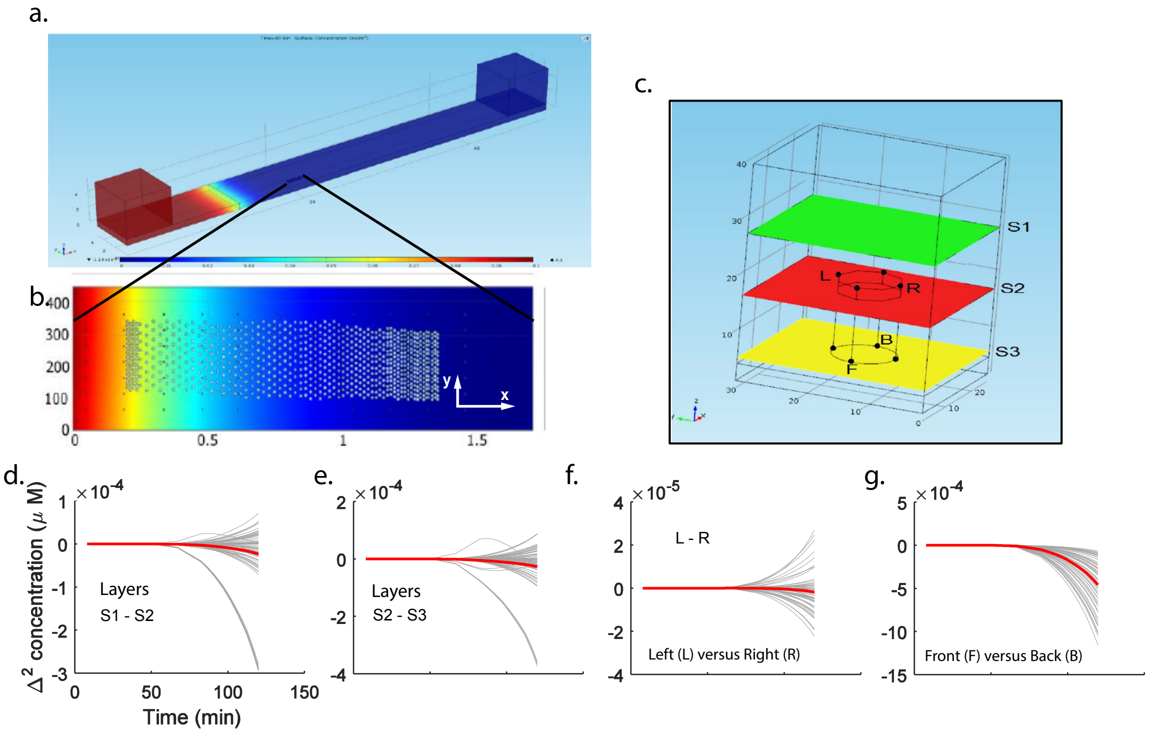


Figure S9: **Modeling the cAMP gradient across the pillar array by finite element modeling a.** Visualization of the cAMP gradient over the pillar array **b.** Visualization of the concentration profile over the array at the end of a measurement at t=150 min. **c.** Investigation of the influence of the pillar geometry on a diffusion gradient by finite element modeling. **d. - e.** The gradient was numerically investigated at three different layers in Z at different positions on the array (grey lines) showing negligble differences in the gradient during the measurement (red lines). **f.** Additionally, the concentration differences between the left (L) and right (R) (y-direction, see **b.**) of many pillars was investigated (grey lines), yielding no significant differences during the experiment (red line). **g.** Finally, the front (F) and back (B) (x-diriction, see **b.**) of many pillars was numerically investigated along the array (grey lines). As expected, the front and back of a pillar generate a consistent chemical gradient with solute differences between 0.01 - 0.5 nM.

[14] David Holcman and Zeev Schuss. *Stochastic Narrow Escape in Molecular and Cellular Biology*. Springer-Verlag New York, 2015.

[15] Zeev Schuss. *Theory and Applications of Stochastic Processes*. Springer-Verlag New York, 2009.

[16] Adolf Fick. On liquid diffusion. *Poggendorffs Annalen*, 94, 1855.
